# A Pilot Study Using Two Novel fMRI Tasks: Understanding Theory of Mind and Emotion Recognition Among Children With ASD

**DOI:** 10.1101/2021.03.24.436877

**Authors:** Yu Han, Patricia A. Prelock, Emily L. Coderre, Joseph M. Orr

**Affiliations:** Department of Communication Sciences and Disorders, University of Vermont, Burlington, VT, United States; Neuroscience Graduate Program, University of Vermont, Burlington, VT, United States; Department of Psychological and Brain Sciences, Texas A&M University, College Station, TX, United States; Texas A&M Institute for Neuroscience, Texas A&M University, College Station, TX, United States

## Abstract

Children with autism spectrum disorder (ASD) struggle with social interactions due to deficits in theory of mind (ToM). In this study, we collected behavioral and neuroimaging data from 9 children with ASD and 19 neurotypical children between the age of 7 and 14 years old, particularly in the area of emotion recognition to better understand those skills needed for meaningful social interaction. The results suggest impaired abilities in multiple ToM metrics and brain deficits associated with ToM-related emotion recognition and processing among children with ASD. Findings from this study are expected to establish connections between behavior and brain activities surrounding ToM in ASD, which may assist the development of neuroanatomical diagnostic criteria and provide a way to measure intervention outcomes.

## Introduction

Autism spectrum disorder (ASD) is a lifelong neurodevelopmental disorder in which an individual’s symptoms can vary from mild to severe. According to the most recent prevalence rates from the US Centers for Disease Control and Prevention (CDC), about 1 in every 54 individuals has ASD (Center for Disease Control and Prevention, 2020). Although many theories exist about the pathology and causes of autism, such as genetic and environmental factors, ASD is a heterogeneous disorder without a specific known cause or cure. Language and intellectual impairments may or may not be characteristic of children with ASD, but the most significant challenges they face are difficulties communicating and interacting with others in social situations (Elsabbagh et al., 2012). Early diagnosis and intervention (e.g., speech and language therapy, social cognitive behavioral intervention, etc.) targeting these social difficulties are especially critical if we wish to improve the social communication skills of children with autism, as well as help them build relationships, engage in activities with others, and be successful in school. An important component of social communication and social interaction in children with ASD is theory of mind (ToM). ToM is the ability to reason about the thoughts and feelings of self and others, including the ability to predict what others will do or how they will feel in a given situation on the basis of their inferred beliefs (Baron-Cohen et al., 1985; Baron-Cohen, 1995). Difficulties with ToM are thought to lead to impairments in social interactions among individuals with ASD. It has been argued that a diminished ability to interpret the beliefs, intentions, and emotions of others will undermine the individual’s ability to interact in ways that are generally considered appropriate and adaptive for a particular social context (Hoogenhout and Smith, 2017). Individuals with ASD often have trouble interpreting or reading the verbal and non-verbal communications of others, specifically in social interactions (American Psychiatric Association, 2020).

ToM abilities have been adopted as proxies to functioning level in ASD for several reasons: (1) the developmentally sequenced acquisition of ToM skills in childhood is well documented (Peterson et al., 2012; Wellman and Liu, 2004); (2) ToM tests have been used in a variety of populations and cultures (Avis and Harris, 1991; Baurain and Grosbois, 2013; Henry et al., 2006); and (3) ToM deficits ostensibly underlie social communication impairments in ASD (Happé and Frith, 1996; Tager-Flusberg, 2007; Volkmar et al., 2014). Additionally, general ToM assessment is internationally applicable in that ToM skills develop in roughly the same manner across the world (Slaughter and Perez-Zapata, 2014; Wellman, Cross, et al., 2011; Wellman, Fuxi, et al., 2011). ToM abilities have also been proposed as a potential severity index in ASD: better ToM is associated with improved behavior towards social rules (Thirion-Marissiaux and Nader-Grosbois, 2007), better social interaction skills (Bosacki and Astington, 1999; Fombonne et al., 1994), and increased language use (Charman et al., 2000; Happé, 1993).

At a neural level, studies have established a ToM network involving the medial prefrontal cortex (mPFC), the posterior superior temporal sulcus (pSTS), the temporal parietal junction (TPJ), the precuneus, and the posterior cingulate cortex (PCC). More specifically, the mPFC is associated with mental state reflection; the pSTS is involved in inferring to other’s actions; and the TPJ with understanding beliefs and socially relevant information (Kana et al., 2015; O’Nions et al., 2014; Takeuchi et al., 2002). Individuals with ASD exhibit decreased activation and connectivity among these identified ToM regions, as well as decreased connectivity in the frontal-medial, frontal-parietal and medial cerebellum anatomical networks (Kana et al., 2015; O’Nions et al., 2014; Takeuchi et al., 2002). The purpose of this study is to examine behavioral and neurobiological measures of emotions involving ToM, contributing to what is known about ToM markers at the brain and behavior levels that can distinguish those with and without ASD. In the review of the literature that follows, we discuss the development of emotion recognition as one aspect of ToM in neurotypical (NT) and ASD populations surrounding happiness, sadness, surprise, embarrassment and desire-based emotion. This includes a description of how emotion recognition has been tested and measured at both a behavioral and neural level in individuals with ASD.

### Emotion Recognition in Neurotypical Development

One particular aspect of ToM, emotion recognition, plays a critical role in an individual’s ability to meaningfully engage in social communication and social interaction. Emotion recognition is the ability to discriminate between different facial expressions and is key to understanding empathy or the feelings of others. The present study focuses on three specific emotions (i.e., surprise, embarrassment, desire-based emotion) as they are critical aspects of ToM.

Happiness is considered to be the easiest recognized emotion while sadness is associated with the most negative affective reactions among the NT population (Mancini et al., 2018). Meta-analyses have found that the processing of emotional faces is associated with increased activation in a number of visual, limbic, TPJ and prefrontal areas, where happy and sad faces specifically also activate the amygdala (Fusar-Poli et al., 2009). Surprise conveys a sense of novelty or unexpectedness and most research indicates that accurate recognition of surprise will happen around the preschool years or even later among the NT population (Weimer et al., 2012). One functional magnetic resonance imaging (fMRI) study suggests that rapid recognition of surprised faces is associated with greater brain activities in the right postcentral gyrus and left posterior insula (Ke et al., 2017).Embarrassment is often described as a ‘self-conscious’ emotion that is associated with a feeling of shame or awkwardness around some action or statement (Castelli, 2005; Hillier and Allinson, 2002a, 2002b; Hutchins and Prelock, 2017). Experiencing ‘embarrassment’ does suggest some level of self-awareness that an ‘expected’ behavior in a social context was unmet (Begeer et al., 2014; Bennett and Matthews, 2000; Edelmann and Hampson, 1981; Edelmann, 1981, 1985; Lewis et al., 1991). Embarrassment is evoked during negative evaluation following norm violations and supported by a fronto–temporo–posterior network. It often recruits greater anterior temporal regions, representing conceptual social knowledge (Jankowski and Takahashi, 2014). Desire-based emotion recognizes the relationship between getting what you want and feeling happy and not getting what you want and feeling sad or disappointed. Thus, desire-based emotion can lead to positive emotions or negative emotions, depending on a fulfilled or unfulfilled desire (Rieffe et al., 2000; Rieffe et al., 2010). There is abundant evidence that around the age of two, NT children understand desire-based emotion and can accurately predict emotional consequences when another’s desire and the situational outcome are known (i.e., others are judged as ‘happy’ if the outcome was wanted and ‘sad’ if it was not) (Ziv and Frye, 2003).

### Emotion Recognition in ASD

Children with ASD have impairments in social interaction often due to a lack of understanding of emotions and the minds of others, as well as difficulty attending to social cues (e.g., gaze, facial expressions, body postures, etc.) (Kuusikko et al., 2009). Some studies have found that children with ASD use the lower part of the face to determine one’s facial expression and often ignore or have difficulty identifying negative facial affects evident near the eyes (e.g., distress, fear) as early as the age of three (Kuusikko et al., 2009). However, other studies suggest that children with ASD have trouble recognizing emotions from the lower part of the face compared to NT children (Kuusikko et al., 2009). There is a wealth of behavioral evidence showing that recognition of even more early developing emotions like happiness and sadness are impaired in individuals with ASD (Apicella et al., 2013; Corbett et al., 2010; Murphy et al., 2012). On the other hand, there is evidence of intact recognition of happiness in some individuals with ASD (Ashwin et al., 2006) as well as a ‘happy advantage’ as recognition of happiness within ASD groups tends to be better than recognition of other emotions (Custrini and Feldman, 1989; Lacroix et al., 2014; Rump et al., 2009; Uljarevic and Hamilton, 2013; Wallace et al., 2008). Better recognition of happiness is also associated with greater social competence (Custrini and Feldman, 1989). The recognition of negative emotions including sadness is generally found to be impaired in ASD (Corden et al., 2008; Kuusikko et al., 2009; Poljac et al., 2012; Wallace et al., 2008). Poor accuracy during sadness recognition tasks is associated with higher symptom severity and poorer adaptive functioning in individuals with ASD (Heerey et al., 2003).

Research has also demonstrated that during face recognition tasks, individuals with ASD show activity in brain areas typically related to the object perception pathway in NT individuals (Koshino et al., 2008; Scherf et al., 2010), suggesting that individuals with ASD may be compensating for a lack of functionality in the core and extended face perception pathways by recruiting regions comprising more general object perception networks. This may explain why ASD individuals perform reasonably well on some behavioral tasks involving emotional face processing (Nomi and Uddin, 2015), perhaps by adopting a compensatory strategy.

The fusiform gyrus (FG), the superior temporal sulcus (STS), and the amygdala have been implicated in the aberrant neuropathology of ASD during face processing. In general, there is evidence for atypical patterns of brain activity in the form of hypoactivation of the FG, STS, amygdala and the occipital lobes, alongside hypoconnectivity of the FG in individuals with ASD. In addition, individuals with ASD demonstrate hypoactivation and hypoconnectivity in areas of the face perception network, including the inferior frontal gyrus (IFG) (Hadjikhani et al., 2007), ITG (inferior temporal gyrus) (Hadjikhani et al., 2007), and middle frontal gyrus (MFG) (Koshino et al., 2008). These results demonstrate that atypical brain activation during emotional face perception is not restricted to the core face perception pathway, but also extends to other cortical areas related to executive functions such as attentional control and inhibition. Taken together, these findings suggest that atypical face perception in ASD is mediated by other factors in addition to pure visual perception.

The neural mechanisms underlying the interpretation of basic emotions such as happy and sad faces in ASD are well studied. Although most studies have reported decreased amygdala activation during emotional face processing (e.g., angry and fearful), one study has found greater right amygdala activation in the ASD group compared to the NT group when processing happy and sad faces (Ashwin et al., 2007; Critchley et al., 2000; Dapretto et al., 2005; Grelotti et al., 2005; Hadjikhani et al., 2007; Pelphrey et al., 2007; Pinkham et al., 2008). Specifically, there was a greater positive functional connectivity between the right amygdala and ventromedial prefrontal cortex to happy faces but less positive functional connectivity between the right amygdala superior/medial temporal gyri (Monk et al., 2010). Other studies also found that the ASD group showed greater bilateral activation in the amygdala, vPFC and striatum comparing to the NT group (Daly et al., 2012). Due to high variability across fMRI studies, other brain regions have also been identified, but overall reduced brain activities when processing happy faces are observed in ASD groups (Monk et al., 2010; Spencer et al., 2011; Weng et al., 2011). The literature has also found consistent results that the ASD population is more sensitive to sad faces. When processing sad faces, ASD groups tend to show greater activation relative to control groups in the amygdala, vPFC, putamen, and striatum, and younger adolescents show greater activation than older adolescents (Daly et al., 2012; Weng et al., 2011). However, one study found decreased activities in mPFC among the ASD group when processing sad faces (Daly et al., 2012).

Contrary to early-developing emotions (e.g., happy, sad, mad, scared) that are responses to situations, recognizing surprise among children with ASD appears to lag behind. Knowing that understanding one’s own and others’ desires, beliefs and values is a particular area of deficit for children with ASD, it is not unexpected they would be challenged in their ability to make sense of the concept of ‘surprise’ (Apicella et al., 2013; Corbett et al., 2010; Hutchins and Prelock, 2017; Murphy et al., 2012). Desire-based emotion plays an important role when understanding and empathizing with others’ thoughts and feelings (Astington, 1993; Nicola, 1984; Peterson and Wellman, 2005; Phillips et al., 1995; Wellman and Woolley, 1990; Ziv and Frye, 2003). In general, the understanding of desire among children with ASD is very limited and they often are unable to generalize their understanding without explicit instruction and support in social contexts (Hutchins and Prelock, 2017). Surprise and desire-based emotions have only been examined at a behavioral level in individuals with ASD and little is known about their neural correlates. Children with ASD may show an emotional response of embarrassment, although to a lesser degree than their NT peers. Further, they seldom recognize embarrassment or express their experiences in situations of embarrassment. Often, they miss social gaffes where what is said is perceived as an inappropriate comment in a social context (Bennett and Matthews, 2000; Burnettand et al., 2009; Hutchins and Prelock, 2017; Modigliani and Blumenfeld, 1979; Seidner et al., 1988; Shamay-Tsoory, 2008; Tangney et al., 1996; Williams and Happé, 2010). Embarrassment has been studied at a neural level only among adults with ASD, with evidence suggesting altered circuitry in the mPFC, ACC, IFG, TPJ/pSTS, posterior cingulate cortex (PCC), and amygdala (Hillier and Allinson, 2002a, 2002b; Jankowski and Takahashi, 2014).

### Purpose of the Study

Although ToM has been studied for decades, it still remains a challenging research area due to its multi-faceted composition. Although the neural mechanisms underlying ToM have been examined, few brain-based studies include children with ASD. Thus, a greater understanding of the brain-behavior connections associated with ToM in children will provide researchers a potential link between the biological mechanisms of ToM and behavioral characteristics. It will help lead to more efficient diagnostic processes and prognostic indicators for special populations like children with ASD. To facilitate increased understanding of this linkage, the current study will emphasize emotion recognition ToM with a specific focus on less well-studied and more complex emotions (i.e., surprise, embarrassment, and desire-based emotion). Surprise and embarrassment are particularly difficult for children with ASD to recognize (Heerey et al., 2003; Uljarevic and Hamilton, 2013). While desire-based emotions are easier for children with ASD to recognize, their understanding of these emotions is delicate and often requires explicit descriptions.

The current study is the first to examine the neural correlates of selected ToM constructs including desire-based emotions and more complex emotions (surprise, embarrassment) to establish connections between behavior and brain activities in children with ASD (i.e., 7 to 14 years old). We used The Theory of Mind Inventory-2 (ToMI-2) (Hutchins and Prelock, 2016) and The Theory of Mind Task Battery (ToMTB) (Hutchins et al., 2008) to establish the behavioral patterns and identify differences between ASD and NT groups in their understanding of these less well-studied and complex emotions. We developed two novel fMRI tasks to identify brain regions associated with the recognition and processing of these emotions. Although our primary interest was in the neural response to recognition of more complex emotions requiring ToM (i.e., embarrassment and surprise), happy and sad faces were also included to provide a comparison with previous literature investigating recognition of basic emotions. We established brain activation patterns in both groups to further probe brain deficits and neural compensation mechanisms being adopted by the ASD group. Specifically, we expected to see altered brain activity patterns among the ASD group in brain regions involved in the ToM neural network (e.g., mPFC, pSTS, cingulate cortex, and TPJ). Building upon findings from previous studies, the present study provides a deeper and more systematic understanding of the brain-behavior connections associated with ToM. This knowledge may lead to both behavioral and neural pathways for examining the impact of intervention research to achieve more normalized social performance. Such research might also help predict those brain-behavior profiles of children most likely to benefit from specific ToM or social cognitive based interventions, emphasizing the importance of interventions delivered at the cognitive level to bridge behavior with brain function (Hutchins and Prelock, 2015).

## Methods

### Participants

Eleven children with ASD (1 female) and 22 NT children (7 females) participated in the study, in which 9 ASD and 19 NT subjects were included as they completed all of the the behavioral testing, magnetic resonance imaging (MRI) scans and fMRI tasks. The remaining subjects either completed only the behavioral testing, or withdrew from the study due to dental appliances precluding MRI scanning or because of study interruption due to COVID-19. The full study included 2-3 hours of baseline behavioral assessments along with a 1-hour brain scan including T1 imaging, T2 imaging, and two fMRI tasks.

Since the understanding of surprise and embarrassment has been shown to develop substantially between the ages of 5 and 8 years in NT populations (Bennett, 1989; Bennett and Matthews, 2000; Bennett et al., 1998), we set the minimum cut off age as 7 years old. We expanded the upper age limit to 14 years old to improve recruitment efforts given the challenges of recruiting subjects with ASD. All children were native English speakers.

We administered the Autism Diagnostic Observation Schedule-2 (ADOS-2) (Lord et al., 1999) and the Social Communication Questionnaire-Lifetime version (SCQ) (Rutter et al., 2003) to confirm the clinical diagnosis for participants with ASD. Non-verbal intelligence and language levels were tested for all participants using the Universal Nonverbal Intelligence Test (UNIT-2) (Bracken and McCallum, 2016; McCallum, 2003) and the Comprehensive Assessment of Spoken Language (CASL), respectively, (Carrow-Woolfolk, 2017) to ensure participants could demonstrate understanding of the instructions given in the behavioral and fMRI tasks. The UNIT-2 is a multidimensional assessment of intelligence for individuals with speech, language, or hearing impairments. It consists of nonverbal tasks that test symbolic memory, non-symbolic quantity, analogic reasoning, spatial memory, numerical series, and cube design. The CASL is an orally administered language assessment consisting of 15 subtests measuring language for individuals ranging from 3 to 21 years of age. For the present study, only those basic subsets that establish the CASL language core were used: Antonyms, Sentence Completion, Syntax Construction, Paragraph Comprehension, and Pragmatic Judgment.

Full demographic statistics are presented in Table 1. The groups differed on CASL and UNIT-2 scores, with the ASD children obtaining lower scores on both measures compared to the NT children. Because of these group differences in language and intellectual abilities, CASL and UNIT-2 scores were included as covariates in statistical analyses of ToM metrics.

**Table 1.** Demographics and T-tests Statistics including Mean, Range and p value

|  | **NT Group**<br>**(n=19)** | **ASD Group**<br>**(n=9)** | **Group**<br>**Difference** |
| --- | --- | --- | --- |
| *Age* | 10.2 (7-14) | 11 (8-13) | $p = 0.36$ |
| *CASL* | 113 (89-132) | 78.8 (40-121) | $p < 0.001^{**}$ |
| *UNIT-2 Full Scale* | 111.9 (100-135) | 96.2 (56-127) | $p < 0.05^{*}$ |
| *ADOS-2* | N/A | 14 (9-21) | N/A |
| *SCQ* | N/A | 22.3 (14-30) | N/A |

### Behavioral Measures of ToM

Two norm-referenced tools were used as behavioral outcome measures to assess ToM. The ToMI-2 (Hutchins and Prelock, 2016) measures a parent’s perception of their child’s ToM understanding of 60 items using a 20-unit rating scale from “Definitely Not” to “Definitely”. Primary caregivers use a vertical hash mark to indicate where on the continuous scale best represents their perceptions. Item, subscale, and composite scores range from 0-20. A higher number indicates a parent’s greater confidence in their child’s understanding of a particular ToM skill. The ToMI-2 items represent typical social interactions to ensure it is a socially and ecologically valid ToM index. The tool demonstrates excellent test-retest reliability, internal consistency, and criterion-related validity for neurotypical children and children with ASD, as well as contrasting-groups validity and statistical evidence of construct validity (i.e., factor analysis) (Hutchins and Prelock, 2016; Hutchins et al., 2012; Lerner et al., 2011).

ToMTB (Hutchins et al., 2008)is a direct measure of a child’s understanding of ToM. It consists of nine ToM tasks presented as short vignettes in a story-book format arranged in ascending difficulty. For each of the nine tasks, children are provided with one correct response option and three possible distracters. There are 15 total questions asked, including memory control questions that must be answered correctly to get credit for ToM understanding. The ToMTB has strong test-retest reliability (Hutchins and Prelock, 2016; Hutchins et al., 2008).

To examine subjects’ ToM abilities, we used the total score of the ToMTB, total composite mean of the ToMI-2 (i.e., assessing overall ToM ability), early subscale mean of the ToMI-2 (i.e., assessing early developing ToMI ability such as regulating desire-based emotion and recognition of happy and sad), basic subscale mean of the ToMI-2 (i.e., assessing basic ToM ability such as recognition of surprise), and advanced subscale mean of the ToMI-2 (assessing advanced ToM ability such as recognition of embarrassment). We also included scores from single items assessing recognition of simple emotions such as happy and sad, as well as more complex emotions such as surprise and embarrassment.

### MRI Acquisition

All neuroimaging data was acquired using the University of Vermont MRI Center for Biomedical Imaging 3T Philips Achieva dStream scanner and 32-channel head coil. The imaging protocol is based on that developed for the multicenter NIH-funded Adolescent Brain Cognitive Development (ABCD) study, which is derived from large studies such as the Human Connectome Project (HCP) and the Lifespan Connectome Project. The protocols make extensive use of simultaneous multislice imaging (Breuer et al., 2005; Setsompop et al., 2012a, 2012b) (multiband SENSE) to accelerate functional and diffusion MRI acquisitions.

### Task fMRI Parameters and Preprocessing

Task fMRI parameters were: TR 800ms, TE 30ms, flip angle 52 degrees, 2.4mm isotropic imaging resolution with a 216×216×144*mm*^3^ field of view using a multiband acceleration factor of 6 (60 slices, no gap). For fMRI acquisitions, corresponding field maps were generated using pairs of reference acquisitions with opposite phase-encode directions. fMRI preprocessing used the pipelines developed as parts of the HCP (Setsompop et al., 2012a). The HCP functional pipeline corrects for EPI spatial distortions using magnetic field maps and realigns volumes to account for subject motion. Specifically, the fMRI surface pipeline was included in the preprocessing analysis while independent component analysis (ICA) denoising was not. Participants were trained to remain still during the scan in a mock scanner supplemented by video model (a short video demonstrating expected behavior in the scan) prior to each assessment. The HCP task fMRI pipelines were used for first- and second-level analysis of task fMRI data. These pipelines incorporate high-pass filtering and application of general linear models (GLMs) to model task parameters and nuisance regressors (e.g., motion parameters) (Jensen et al., 2005; Jeurissen et al., 2014).

### fMRI Emotion Recognition (fER) Task

This task was designed to assess emotion recognition, which was also a behavioral item tested on both ToMI-2 and ToMTB (Hutchins and Prelock, 2016; Lerner et al., 2011). The fMRI task required participants to either identify the emotions expressed in cartoon faces (emotion recognition) or judge the gender of the same emotional faces (perceptual control). Faces depicting happiness, sadness, surprise, and embarrassment were included. Since we did not find existing stimulus/picture sets depicting embarrassment or of facial expressions of children in general, we hired a professional artist to create digital drawings of the cartoon face stimuli that matched the age of the participants and the style of pictures used in the ToMTB and ToMI-2. An independent sample of 163 NT participants from Amazon Mturk validated and selected the expressions that were recognized with 80% to 90% accuracy and were matched for valence. The final selection included eight different characters and two versions of each expression for each character, leading to 16 unique stimuli for each expression (see Figure 1 panel b for examples).

**Figure 1.**
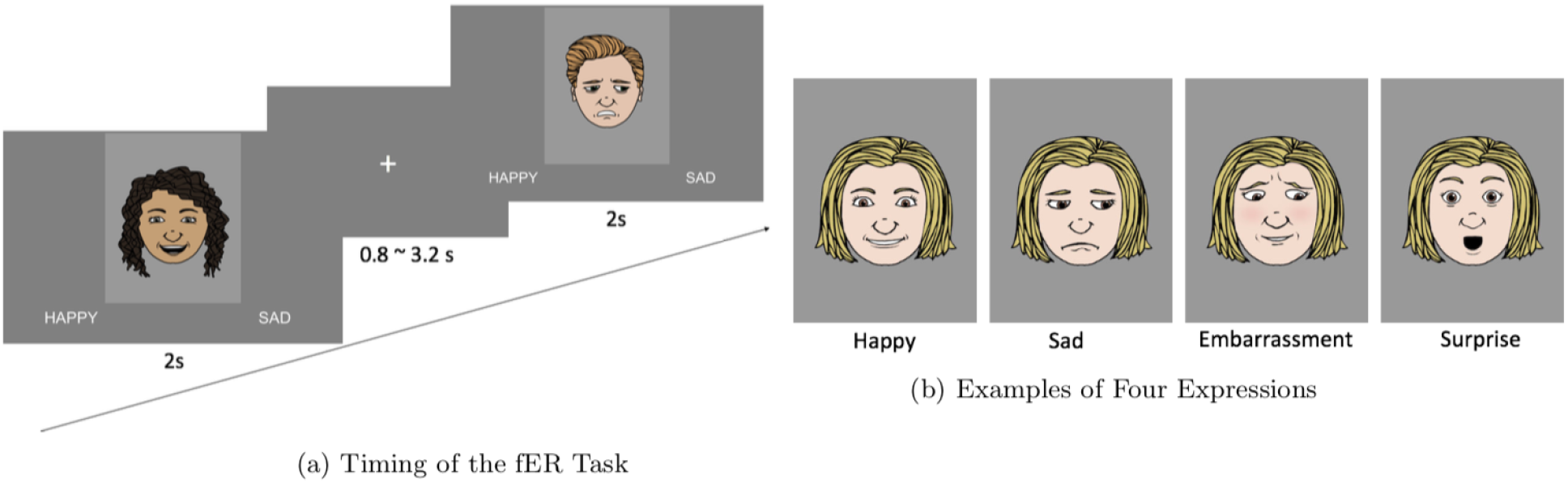
Illustration of the fMRI Emotion Recognition (fER) Task

The fER task utilized a mixed design, with a block design for task (emotion recognition of surprise/embarrassment, emotion recognition of happy/sad, and perceptual control) and an event-related design for facial expression within each block (surprise/embarrassment or happy/sad). Across two runs, we presented 8 blocks of emotion recognition of surprise and embarrassment, 8 blocks of emotion recognition of happy and sad, and 8 blocks of perceptual control. Each block presented 8 faces in an event-related fashion for 2 seconds each with a jittered ISI (i.e., multiples of the TR, optimized using the optseq tool (FreeSurfer, 2012). Participants pressed one of two buttons to indicate the emotional expression (surprise/embarrassment or happy/sad, in emotion recognition blocks) or the gender of the face (boy/girl, in perceptual control blocks; see Figure 1, panel a). An instructional cue (i.e., Label Emotion, Label Gender) was provided before each block. Between blocks there was an 8 second interval. The mixed design allowed us to not only create different contrasts between each emotion, but also between advanced emotion processing (i.e., recognition of combined embarrassment and surprise) with basic emotion processing (i.e., combined happy and sad).

### fMRI Theory of Mind (fToM) Task

We developed this fMRI task to directly model the desire-based emotion task in the ToMTB (Hutchins et al., 2008). Participants were required to infer the reaction of a cartoon character to a gift using the knowledge provided about the preferences of that character. Each trial was broken into three images: encoding, probe, and decision-making (see Figure 2). In the encoding image (2.4s), a child was seen holding a gift box. Two items were presented in thought bubbles displaying the preferences for that character. One item was presented with hearts to indicate the character likes the item, the other with an X over it to indicate the character dislikes that item. In the probe image (2s), a top view of the unwrapped gift box showed the desired/undesired item. Finally, in the decision-making image (2s), participants saw two images of the character’s face expressing either happiness or sadness and respond by pressing a button to indicate which face best captured the reaction of the character to the gift. In control trials, participants saw an empty thought bubble and gift box and were asked at the decision-making step to indicate the gender of the cartoon face. There were 10 different characters and 30 unique items ranging from items commonly recognized as desired by children (e.g., lollipop) to items commonly recognized as unwanted (e.g., spider). This task was presented in an event-related manner over 2 runs and included 15 trials where the character got what he/she likes, 15 trials where the character did not get what he/she likes, and 30 control trials, with a jittered inter-trial interval (ITI) of 0.8-3.2s.

**Figure 2.**
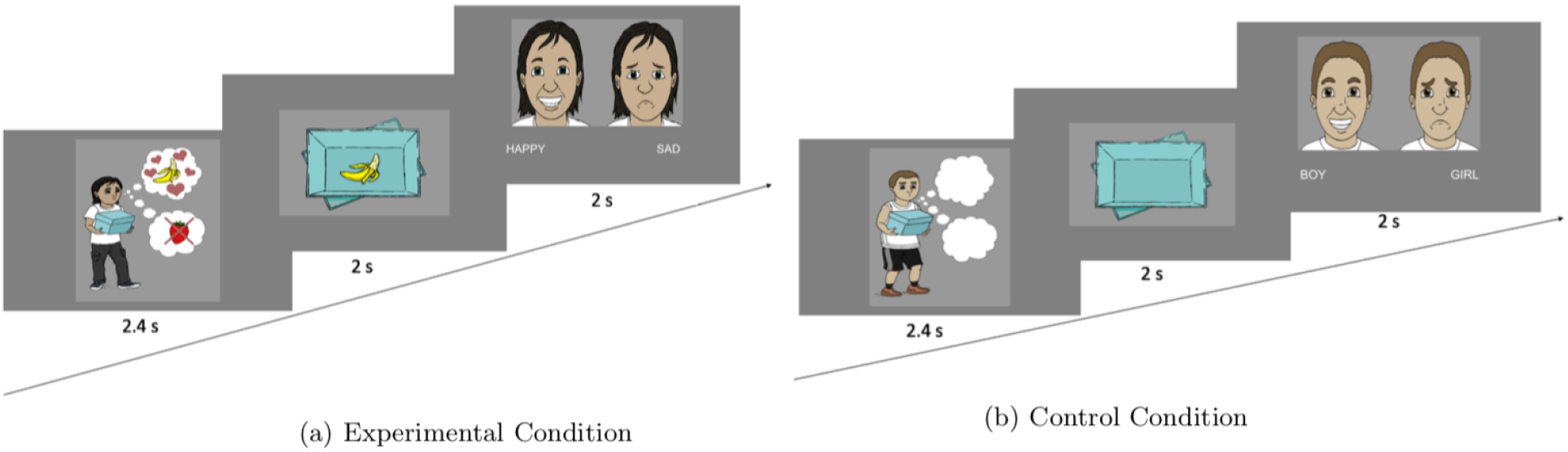
Illustration of the fMRI Theory of Mind (fToM) Task

### Statistical Analyses

Behavioral analyses compared the ToM metrics from the ToMI-2 and the ToMTB, as well as the response time and accuracy from the two fMRI tasks. For analyses of the ToMI-2 and the ToMTB, group comparisons were performed using an analysis of covariance (ANCOVA) with group as a between-subjects factor and CASL score (i.e., language assessment) and UNIT-2 score (i.e., non-verbal intelligence assessment) as covariates. For the fER task, behavioral response times and response accuracy were evaluated for both NT and ASD groups using independent sample *t*-tests for: all conditions combined (overall), surprise condition, embarrassment condition, happy condition, sad condition, happy and sad conditions combined (HS, to examine basic emotion processing), embarrassment and surprise condition combined (ES, to examine advanced emotion processing), and the control (gender) condition. For the fToM task, response times and response accuracy were evaluated for both NT and ASD groups using independent sample *t*-tests for: all conditions combined (overall), control (gender) condition, and experimental conditions (dislike and like conditions combined).

In order to detect neural activation related to the two fMRI tasks, first-level neuroimaging analyses were performed first. Vectors of stimuli onsets using the stimulus duration were created for each trial type and were convolved with a canonical double gamma hemodynamic response function to produce a regressor for each condition. Six motion regressors were included as covariates. For second-level neuroimaging group analyses, whole-brain between-group changes in activation patterns over time were assessed for each experimental condition using FSL’s permutation-based non-parametric testing and threshold-free cluster enhancement to control for multiple comparisons (Smith and Nichols, 2009). Analyses were restricted to gray matter. Different group GLMs were constructed for each hypothesis separately. For the fER task, GLM models were constructed for contrasts between happy vs. gender, sad vs. gender, surprise vs. gender, embarrassment vs. gender, happy & sad vs. gender (HS, to assess basic emotion processing), and embarrassment & surprise vs. gender (ES, to assess advanced emotion processing). For fToM task, GLM models were constructed for contrasts between dislike & like (experimental) vs. control. CASL scores (i.e., language assessment) and UNIT-2 scores (i.e., non-verbal intelligence assessment) were also included as covariates in the permutation analysis of linear models (PALM) for group comparisons (Winkler et al., 2014).

## Results

### Behavioral Results

When adjusted for covariates of the CASL (i.e., language assessment) and UNIT-2 scores (i.e., non-verbal intelligence assessment), the NT group had a significantly better performance on the ToMI-2 total score (*F* (1,24)=4.48, *p*<0.05) and the ToMI-2 early subscale score (*F* (1,24)=7.76, *p*<0.01; see Table 2).

**Table 2.** ToM Behavioral Measurements and ANCOVA Results

|  |  | **NT Group**<br>**(n=19)** | **ASD Group**<br>**(n=9)** | **Group**<br>**Difference** |
| --- | --- | --- | --- | --- |
| **ToMTB** |  | **13.3(11-15)** | **11.1 (5-14)** | **$p = 0.38$** |
| **ToMI-2** | **Total** | **17.2 (11.5-19.3)** | **12.4 (9.1-17.6)** | **$p < 0.05^*$** |
|  | **Early Subscale** | **17.7 (13.8-20)** | **14.6 (11.6-17.9)** | **$p < 0.01^*$** |
|  | **Basic Subscale** | **18 (13.2-19.7)** | **14 (8.5-19.2)** | **$p = 0.12$** |
|  | **Advanced Subscale** | **16 (8.3-18.9)** | **9.4 (6-16.4)** | **$p = 0.07$** |
|  | **Happy** | **18.5 (14.1-20)** | **17.3 (12.5-20)** | **$p = 0.25$** |
|  | **Sad** | **17.5 (7.1-20)** | **16.9 (13.1-20)** | **$p = 0.66$** |
|  | **Surprise** | **17.8 (12.2-20)** | **15.3 (10.3-20)** | **$p = 0.07$** |
|  | **Embarrassment** | **15.3 (5.8-20)** | **10.3 (4.8-20)** | **$p = 0.11$** |
|  | **Desire-Based** | **18.9 (15-20)** | **16.7 (14.6-20)** | **$p = 0.09$** |

On the fER task, the NT group responded significantly faster than the ASD group when considering all conditions together (*t*(54)=2.56, *p*=0.01; see Figure 3), as well as in conditions of surprise (*t*(54)=2.58, *p*=0.01), embarrassment (*t*(54)=2.64, *p*=0.01), sadness (*t*(54)=2.03, *p*<0.05), control (*t*(54)=2.73, *p*<0.01), and ES (embarrassed + surprise) (*t*(54)=2.77, *p*<0.01). There was no significant difference found for the happy condition (*t*(54)=1.75, *p*=0.09) or the HS (happy + sad) condition (*t*(54)=1.95, *p*<0.06). On this task the NT group also had a significantly higher response accuracy compared to the ASD group when considering all conditions together (*t*(54)=2.55, *p*=0.01), as well as in conditions of ES (*t*(54)=2.29, *p*=0.03) and control (*t*(54)=2.52, *p*=0.02). There was no significant difference in terms of response accuracy between the two groups in conditions of surprise (*t*(54)=1.92, *p*=0.06), embarrassment (*t*(54)=1.87, *p*=0.07), happiness (*t*(54)=0.77, *p*=0.44), sadness (*t*(54)=1.89, *p*=0.06), or HS (*t*(54)=1.77, *p*=0.08).

**Figure 3.**
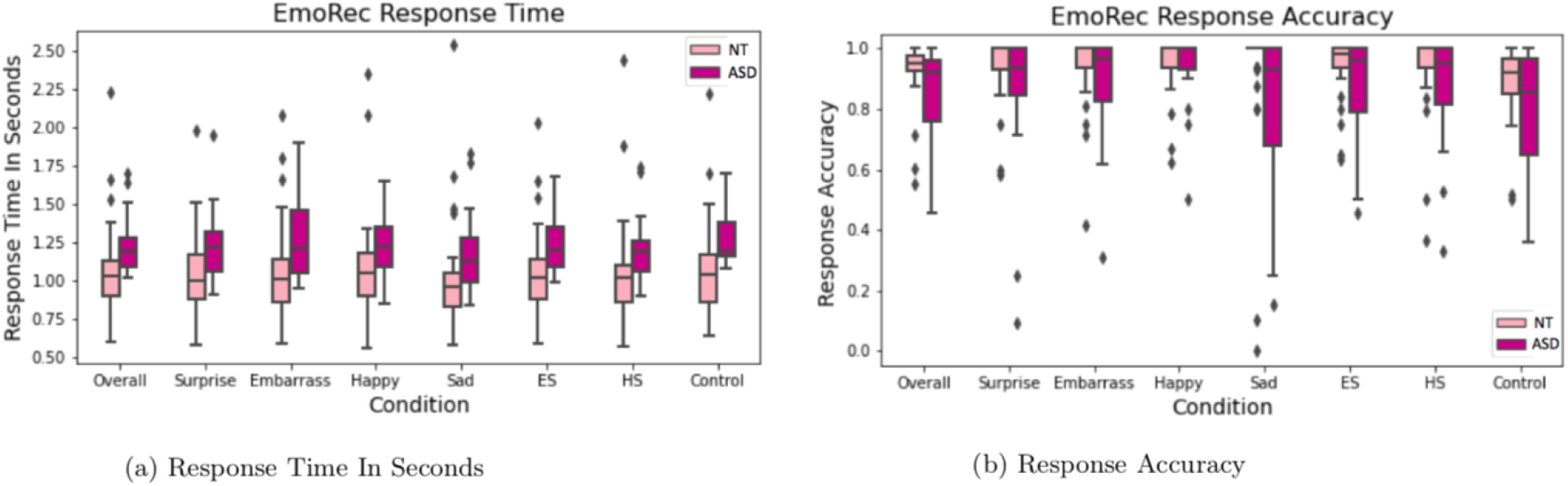
fMRI Emotion Recognition (fER) Task Response Time and Accuracy

On the fToM task, there was no significant difference in response time between the NT and ASD groups for any of the comparisons: all conditions combined (*t*(54)=1.27, *p*=0.21; see Figure 4), experimental condition (*t*(54)=1.14, *p*=0.26), and control condition (*t*(54)=1.33, *p*=0.19). The NT group had a significantly higher response accuracy compared to the ASD group when considering all conditions combined (*t*(54)=2.21, *p*=0.03) and for the experimental condition only (*t*(54)=2.23, *p*=0.03). There was no significant difference in response accuracy for the control condition (*t*(54)=1.92, *p*=0.06).

**Figure 4.**
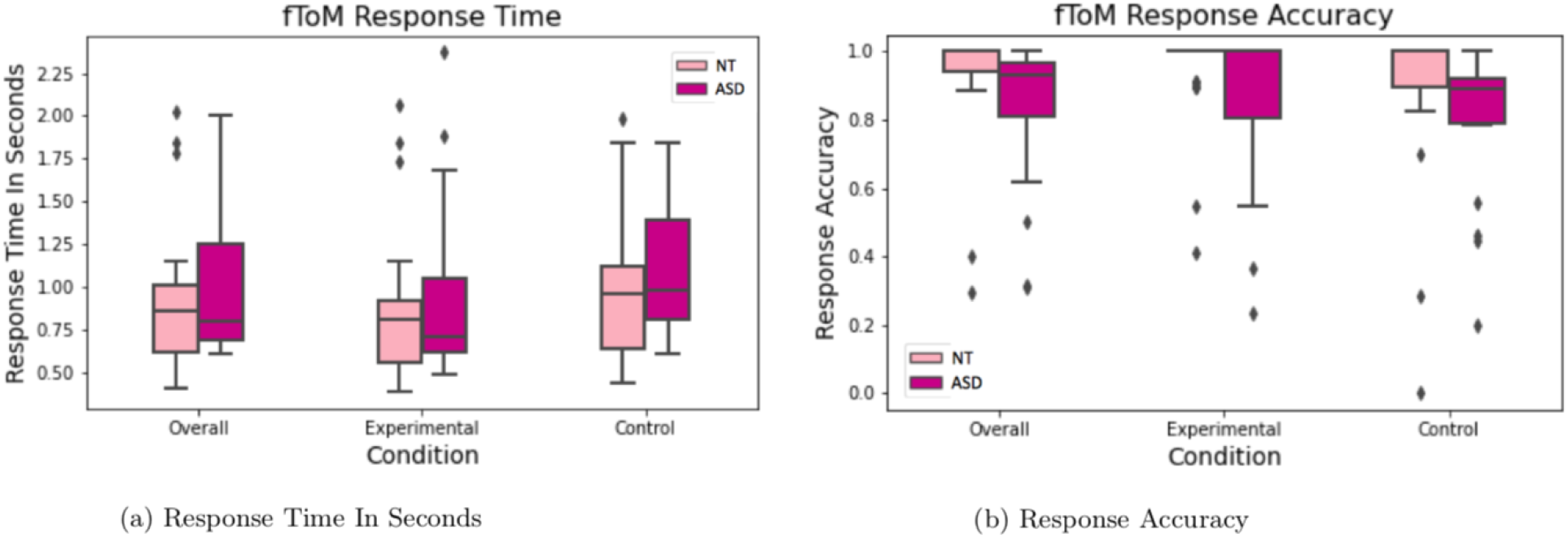
fMRI Theory of Mind (fToM) Task Response Time and Accuracy

### Brain Activity Patterns

Cohen’s D effect size maps were generated for each contrast of both fMRI tasks. The present study adopted a threshold of *d*=0.5 (medium) to show the effect size of the difference in brain activities between the ASD and NT groups. The contrast was conducted as ASD minus NT. Positive d-values (hot colors) indicate greater activation in the ASD group compared to the NT groups, while negative d-values (cool colors) indicate less activation in the ASD group compared to the NT groups.

In the fER task, when recognizing happy faces the ASD group showed greater brain activation in the mPFC and the angular gyrus (AG) compared to the NT group (see Figure 5), but less brain activation in most areas of the frontal cortex, the temporal lobe especially around the inferior temporal sulcus (ITS), and the TPJ. When recognizing sad faces, the ASD group showed greater brain activation in the left temporal lobe, the left AG, the anterior and posterior cingulate cortex, the occipital lobe, and the perirhinal area, but less brain activation in the right temporal lobe, the mPFC, and the inferior part of the post central gyrus. When combining happy and sad faces (i.e., HS), the brain activation pattern was similar to recognizing sad faces alone.

**Figure 5.**
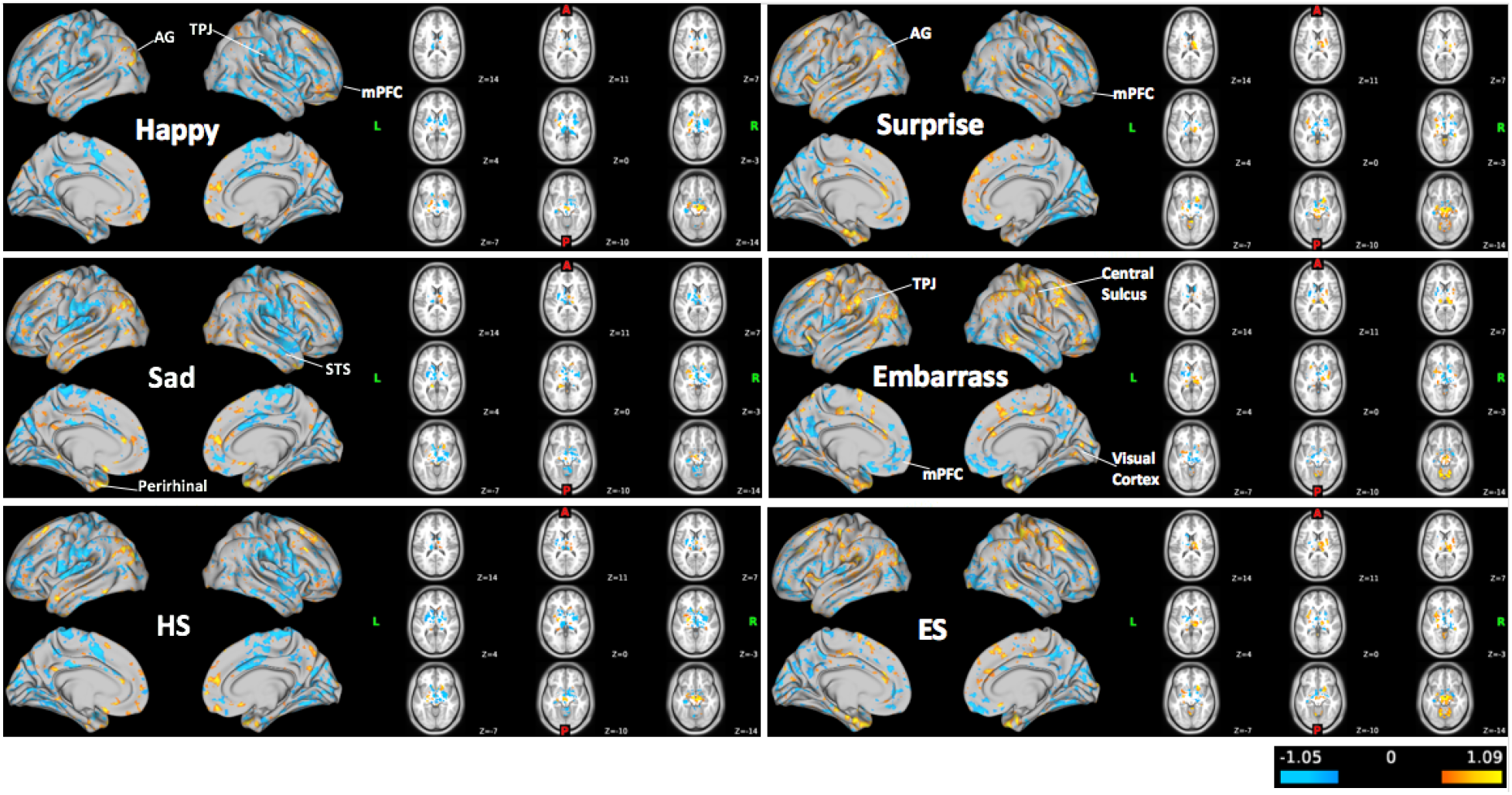
fMRI Emotion Recognition (fER) Task Brain Activation CohenD Maps, thresholded at d=0.5

When recognizing surprised faces in the fER task, the ASD group showed greater brain activation in the left AG, the left temporal pole and STS, the left anterior and posterior cingulate cortex, and the perirhinal area, but less brain activities in the visual cortex and the mPFC, comparing to the NT group (see Figure 5). When recognizing embarrassed faces, the ASD group showed greater brain activities in the TPJ, the AG and the right pre and post central gyrus, the visual cortex and the perirhinal area, but less brain activities in the mPFC. When combining surprise and embarrassed faces (i.e., ES), the brain activation pattern was similar to recognizing embarrassment faces alone.

In the fToM task examining desire-based emotions, the ASD group showed greater brain activation in most of the frontal regions especially around the right mPFC, the AG, the cingulate cortex, and the posterior STS, but less brain activation in the left TPJ and the left temporal lobe, comparing to the NT group (see Figure 6).

**Figure 6.**
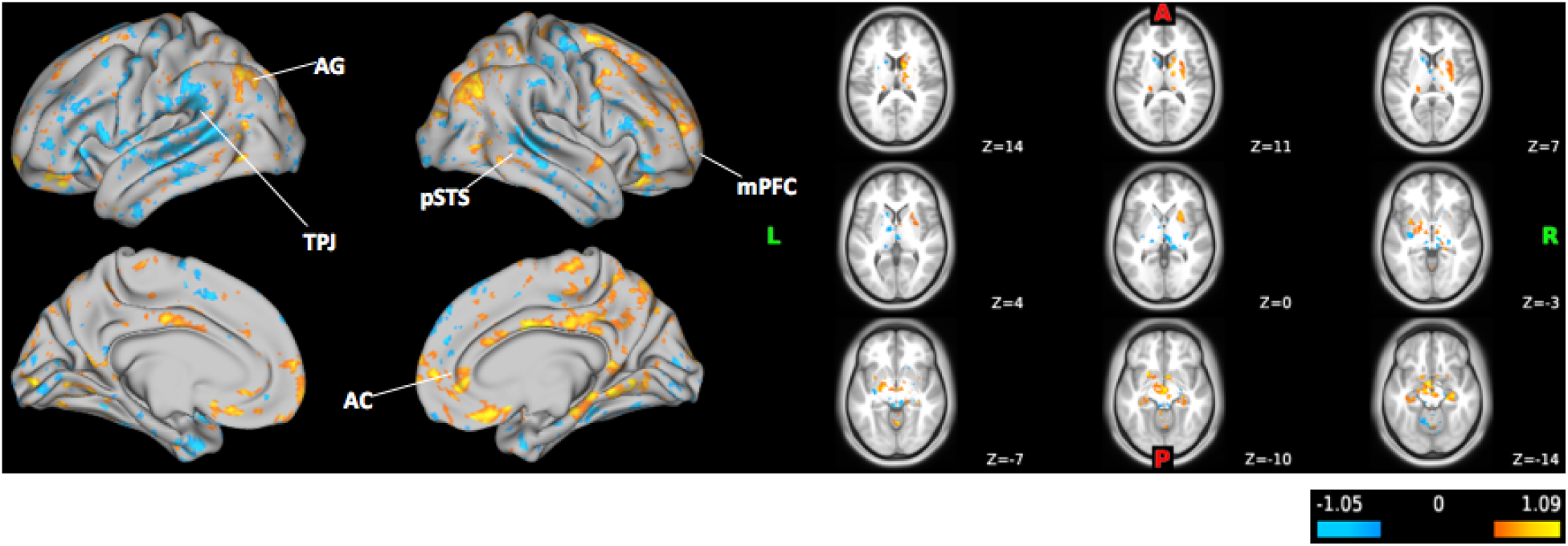
fMRI Theory of Mind (fToM) Task Brain Activation CohenD Maps, thresholded at d=0.5

## Discussion

The current study adopts behavioral and neuroimaging measurements to examine the brain-behavior connections of ToM among children with ASD, specifically in the area of emotion recognition. The results demonstrate behavioral impairments and different brain activation patterns when performing ToM-related tasks in the ASD group compared to the NT group. Scores of behavioral tests suggest that the ASD group has poorer abilities and skills in multiple ToM metrics assessed by the ToMI-2 (Hutchins and Prelock, 2016) and ToMTB (Hutchins et al., 2008) before taking language and non-verbal intelligence levels into account. Specifically, as predicted, the ASD group has more difficulty in recognizing and processing surprise, embarrassment and desire-based emotions, but are equally as good as NT participants at recognizing and processing happy and sad emotions. However, when language and non-verbal intelligence levels are considered, the group differences are only apparent in the ToMI-2 total and early subscale domains. These findings suggest that impairments of ToM abilities are largely associated with language and intellectual levels, especially during more complex emotion recognition. In addition, these findings are consistent with the diagnostic criteria and other common challenges in ASD, including language and intellectual impairments and social challenges requiring advanced ToM abilities.

According to the results from the fER task, the ASD group takes longer to recognize facial expressions of surprise, embarrassment and sadness. This phenomenon is especially apparent when combining conditions of surprise and embarrassment (ES). This is consistent with previous studies suggesting intact ability for recognizing happiness in children with ASD (Ashwin et al., 2006) but impairments in recognizing sadness (Ashwin et al., 2006; Corden et al., 2008; Poljac et al., 2012; Wallace et al., 2008). There are more severe impairments among children with ASD for recognizing surprise and embarrassment compared to happiness and sadness, as these advanced emotions require more cognitive processes (Heerey et al., 2003; Uljarevic and Hamilton, 2013). Thus, it makes sense that the ASD group needs more time to recognize embarrassment and surprise faces. In addition, the ASD group does not recognize surprise and embarrassment faces as accurately as the NT group. Results from the fToM task also indicate poorer performance (i.e., response accuracy) in processing desire-based emotion in the ASD group than the NT group, providing evidence that children with ASD have impaired skills in understanding ToM-associated desire-based emotion.

The brain activation pattern generated by the novel fMRI tasks demonstrate some consistent findings from previous literature. Specifically, when recognizing basic emotions such as happy and sad, the ASD group displays unusual brain activation patterns in the mPFC, the temporal sulcus, and the AG, compared to the NT group. In addition, compared to happiness, recognition of sadness in the ASD group seems to engage a more diverse range of brain regions including the occipital lobe, the left temporal lobe and the perirhinal area associated with the amygdala. This implies that the ASD population has more difficulty recognizing sadness compared to happiness as there is greater brain effort as well as different brain areas that are recruited as compensatory mechanisms (Ashwin et al., 2006; Corden et al., 2008; Custrini and Feldman, 1989; Heerey et al., 2003; Lacroix et al., 2014; Poljac et al., 2012; Rump et al., 2009; Uljarevic and Hamilton, 2013; Wallace et al., 2008). On the other hand, when recognizing happiness there is less brain activation in the TPJ but more brain activation in the AG among the ASD group as compared to the NT group. Multiple studies have established the collaborative relationship between the TPJ and AG during ToM-related tasks among children with ASD, where the AG would display significantly increased activation if a particular ToM task required TPJ activation (Ilzarbe et al., 2020; Schurz et al., 2017). When recognizing more complex emotions such as surprise and embarrassment, the ASD group displayed more brain activation in the left AG but less brain activity in the mPFC area compared to the NT group. This suggests that advanced emotion recognition requires more ToM related abilities than abilities related to executive functions among the ASD group. It also seems that ToM requires more effort for the ASD group to recognize embarrassment compared to surprise as the ASD group engages in more brain activation in regions of the TPJ, the pre and post central gyrus, the visual cortex and the perirhinal area. This implies that the recognition of embarrassment requires more visual information processing and advanced ToM abilities, and may trigger the brain’s fear center (Bastin et al., 2016; Geng et al., 2018; Müller-Pinzler et al., 2015). When processing desire-based emotion, the ASD group displays more brain activity around the right mPFC, the AG, the anterior cingulate cortex, and the posterior STS, but less brain activity around the left TPJ and the left temporal lobe, compared to the NT group. It is obvious that desire-based emotion processing is associated with traditional ToM related brain network including the TPJ, the STS, and the cingulate cortex. With decreased brain activation in the TPJ area, the ASD group seems to engage in the AG as a compensatory mechanism. In addition, the ASD group showed increased brain activity in the mPFC area compared to the NT group suggesting the possibility of recruiting more executive control regions to process the task to compensate for poorer ToM abilities.

As a pilot study, one major contribution of the present study is the successful implementation of two novel fMRI tasks targeting emotion recognition and processing associated with ToM. To our knowledge, this is the first time a set of facial stimuli were able to be adapted to an fMRI task to include expressions of surprise and embarrassment using the faces of children. It is also the first time that neural mechanisms underlying desire-based emotion processing are examined through fMRI. Further, these tasks are directly adapted from the ToMTB and ToMI-2 which allows comparison between behavioral and neuroimaging measurements. In addition, nearly all behavioral tests of ToM currently available only examine one or a few aspects of ToM; however, the ToMI-2 and ToMTB used in the present study are multi-faceted tools that cover several aspects of ToM (e.g., emotion recognition, false belief, perspective taking etc.), including information from both a parent and the child. These tools have helped to identify specific domains in ToM that are a struggle for ASD children. The present study has also identified brain activation maps associated with such domains by using two novel fMRI tasks. These maps provide evidence of how different brain regions are involved in the ability to reflect on mental states (i.e., mPFC), understand other’s actions (i.e., pSTS), integrate relevant social information (i.e., TPJ), create and process emotions (i.e., cingulate cortex), and regulate negative emotions (i.e., amygdala) (Daly et al., 2012; Kana et al., 2015; O’Nions et al., 2014; Takeuchi et al., 2002; Tanaka and Sung, 2017).

## Limitations

Studies using MRI/fMRI technology face the typical challenges for any population with the requirements for tolerating loud noises, remaining still during the assessment and managing feelings of claustrophobia. For an ASD population these challenges are exacerbated, particularly in the recruitment phase of the study and in obtaining usable data. These challenges along with children being fitted with dental pieces precluded our ability to complete the MRI/fMRI images and COVID restrictions further limited access to an ASD population.

To address the challenges for using MRI/fMRI, all participants participated in a mock scanner experience. They viewed two videos explaining the process including one video specifically made to rehearse the experience. These strategies supported the successful participation of many of the participants with ASD but not all, which led to missing data. As a part of our recruitment efforts, we expanded the age limit to 14 years old. The age of the participants in this study ranged from early childhood to the early adolescent period, typically an ideal age range to measure brain size while considering other possible neuroanatomical variables. Importantly, as the brain continues to grow and change into adulthood with rapid and significant development among adolescence (Konrad et al., 2013; National Institute of Mental Health, 2020), the interpretation of the results provides insights for children in general, but does not provide age-specific information. Further, language abilities varied, so the possibility exists that participants may not have fully comprehended the instructions of the fMRI tasks and hence given wrong or negligent responses. To address these concerns, all participants were provided with in-person introductions for both tasks which were also practiced on a computer with the same response device outside the scanner to ensure correct understanding and performance. During the scan, a research team member monitored the response signal box in the monitor room at all times to reassure that participants were paying attention to the task and following task instructions.

## Conclusions and Implications

Currently, the diagnosis of ASD is based on behavioral symptoms alone. Delays in diagnosis of ASD can be 13 months and longer for minority and lower socioeconomic status groups (Bernier et al., 2010; Centers for Disease Control and Prevention, 2012; Mandell et al., 2002; Morrier et al., 2008; Wiggins et al., 2006). It is also believed that a substantial number of individuals on the spectrum remain undetected (Thabtah and Peebles, 2019). Children with ASD are a challenging population to engage in experimental procedures that have specific task requirements and involve neuroimaging measurements. However, it is crucial to gather both behavioral and neuroimaging data to increase our understanding of the nature of the social deficits characteristic in ASD. This is important if we wish to advance the diagnostic process and implement intervention methods for which we are able to show positive behavioral and neural outcomes. Insights derived from this study are expected to help scientists and physicians understand how social deficits surrounding ToM in ASD are associated with the brain. By taking advantage of such knowledge, we might be able to offer insights into the development of neuroanatomical diagnostic criteria to allow for more efficient diagnosis and early identification of high-risk populations. It might also help provide a vehicle for examining neural and behavioral outcomes following individualized treatment methods. Although the present study has some limitations including a relatively small sample size, the successful implementation of two novel fMRI tasks among children with ASD and the comprehensive findings from both behavioral and neuroimaging perspectives should stimulate future research when working with special populations such as those with ASD.

## Supplemental Materials

**Figure 7.**
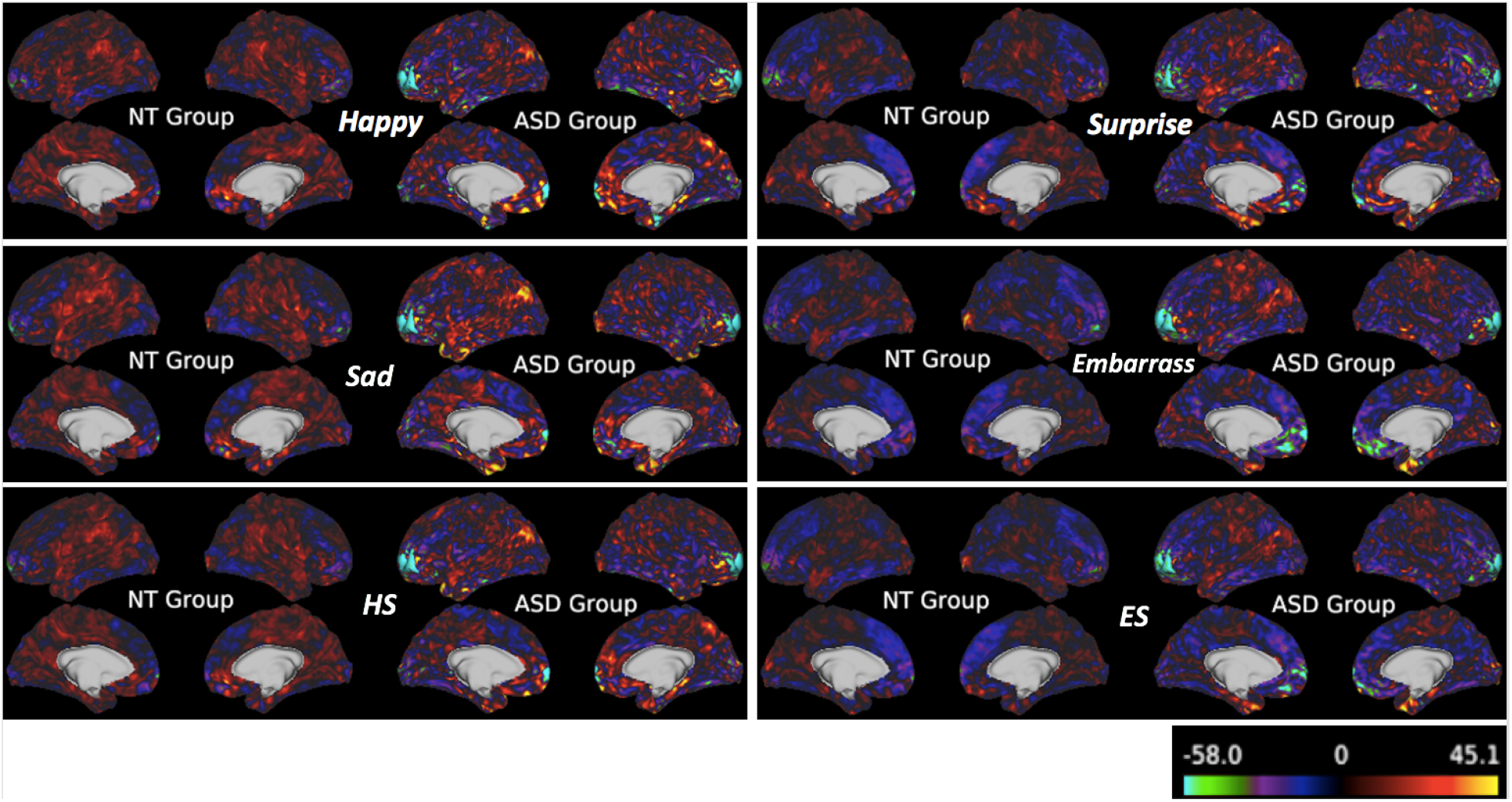
fMRI Emotion Recognition (fER) Task Brain Activation Beta Maps. Beta maps corresponding to voxel-wise mean beta values calculated from the GLM model for each group show the general activation patterns for each contrast. ASD group shows apparent decreased brain activities in the frontal pole region for all conditions and increased brain activities in the angular gyrus and the mPFC when recognizing happy and sad faces.

**Figure 8.**
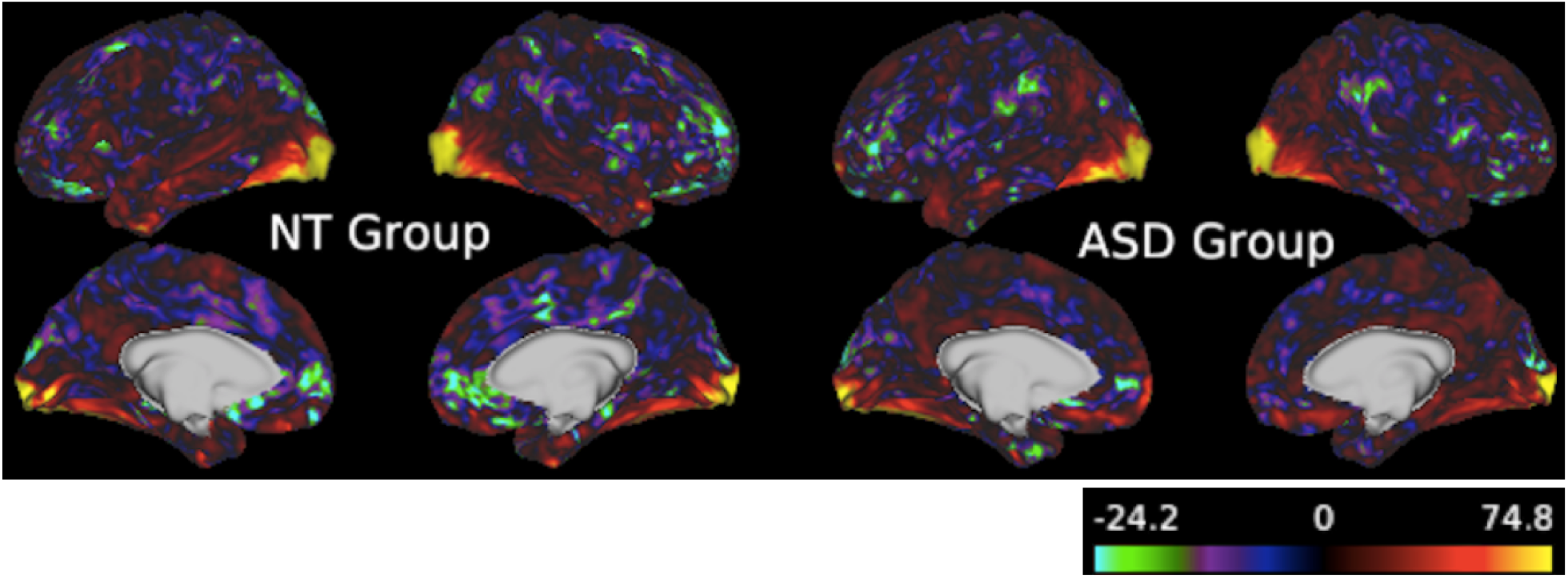
fMRI Theory of Mind (fToM) Task Brain Activation Beta Maps. Beta maps corresponding to voxel-wise mean beta values calculated from the GLM model for each group show the general activation patterns. Both groups show apparent increased brain activities in the visual cortex.

## References

American Psychiatric Association. (2020). Diagnostic and statistical manual of mental disorders (dsm–5).

Apicella, F. et al.. (2013). Fusiform gyrus responses to neutral and emotional faces in children with autism spectrum disorders: A high density erp study. Behavioral Brain Research.

Ashwin, C. et al.. (2007). Differential activation of the amygdala and the ‘social brain’ during fearful face-processing in asperger syndrome. Neuropsychologia.

Ashwin, C., Chapman, E., Colle, L., & Baron-Cohen, S. (2006). Impaired recognition of negative basic emotions in autism: A test of the amygdala theory. Soc Neurosci.

Astington, J. (1993). The childs discovery of the mind. Harvard University Press.

Avis, J., & Harris, P. (1991). Belief-desire reasoning among baka children: Evidence for a universal conception of mind. Child Dev.

Baron-Cohen, S. et al.. (1985). Does the autistic child have a “theory of mind”? Cognition.

Baron-Cohen, S. (1995). Learning, development, and conceptual change. mindblindness: An essay on autism and theory of mind. The MIT Press.

Bastin, C. et al.. (2016). Feelings of shame, embarrassment and guilt and their neural correlates: A systematic review. Neurosci Biobehav Rev.

Baurain, C., & Grosbois, N. N. (2013). Theory of mind, socio emotional problem solving, socio-emotional regulation in children with intellectual disability and in typically developing children. J Autism Dev Disord.

Begeer, S. et al.. (2014). Understanding of emotions based on counterfactual reasoning in children with autism spectrum disorders. Autism.

Bennett, M. (1989). Childrens self-attribution of embarrassment. British Journal of Developmental Psychology.

Bennett, M., & Matthews, L. (2000). The role of second-order belief-understanding and social context in childrens self-attribution of social emotions. Social Development.

Bennett, M., Yuill, N., Banerjee, R., & Thomson, S. (1998). Childrens understanding of extended identity. Developmental Psychology.

Bernier, R., Mao, A., & Yene, J. (2010). Psychopathology, families and culture: Autism. Child Adolesc Psychiatr Clin.

Bosacki, S., & Astington, J. (1999). Theory of mind in preadolescence: Relations between social understanding and social competence. Social Development.

Bracken, B., & McCallum, R. (2016). Universal nonverbal intelligence test 2nd edition. Pro-ed.

Breuer, F. et al.. (2005). Controlled aliasing in parallel imaging results in higher acceleration (caipirinha) for multi-slice imaging. Magnetic Resonance in Medicine.

Burnettand, S. et al.. (2009). Development during adolescence of the neural processing of social emotion. Journal of Cognitive Neuroscience.

Carrow-Woolfolk, E. (2017). Comprehensive assessment of spoken language 2nd edition. Pearson.

Castelli, F. (2005). Understanding emotions from standardized facial expressions in autism and normal development. Autism.

Center for Disease Control and Prevention. (2020). Community report on autism.

Centers for Disease Control and Prevention. (2012). Prevalence of autism spectrum disorders–autism and developmental disabilities monitoring network, 14 sites, united states, 2008. MMWR Surveill Summ.

Charman, T. et al.. (2000). Testing joint attention, imitation, and play as infancy precursors to language and theory of mind. Cognitive Development.

Corbett, B. A. et al.. (2010). A functional and structural study of emotion and face processing in children with autism. Psychiatry Res.

Corden, B., Chilvers, R., & Skuse, D. (2008). Avoidance of emotionally arousing stimuli predicts social-perceptual impairment in asperger’s syndrome. Neuropsychologia.

Critchley, H. et al.. (2000). The functional neuroanatomy of social behaviour: Changes in cerebral blood flow when people with autistic disorder process facial expressions. Brain.

Custrini, R., & Feldman, R. (1989). Children’s social competence and nonverbal encoding and decoding of emotions. Journal of Clinical Child Psychology.

Daly, E. et al.. (2012). Serotonin and the neural processing of facial emotions in adults with autism. an fmri study using acute tryptophan depletion. Arch Gen Psychiatry.

Dapretto, M. et al.. (2005). Understanding emotions in others: Mirror neuron dysfunction in children with autism spectrum disorders. nature neuroscience.

Edelmann, R., & Hampson, S. (1981). The recognition of embarrassment. Personality and Social Psychology Bulletin.

Edelmann, R. (1981). Embarrassment: The state of the research. Current Psychological Reviews.

Edelmann, R. (1985). Social embarrassment: An analysis of the process. Journal of Social and Personal Relationships.

Elsabbagh, M. et al.. (2012). Global prevalence of autism and other pervasive developmental disorders. Autism Res.

Fombonne, E. et al.. (1994). Adaptive behaviour and theory of mind in autism. European Child & Adolescent Psychiatry.

FreeSurfer. (2012). Optseq2. http://surfer.nmr.mgh.harvard.edu/optseq.

Fusar-Poli, P. et al.. (2009). Functional atlas of emotional faces processing: A voxel-based meta-analysis of 105 functional magnetic resonance imaging studies. J Psychiatry Neurosci.

Geng, Y. et al.. (2018). Oxytocin facilitates empathic- and self-embarrassment ratings by attenuating amygdala and anterior insula responses. Front. Endocrinol.

Grelotti, D. et al.. (2005). Fmri activation of the fusiform gyrus and amygdala to cartoon characters but not to faces in a boy with autism. Neuropsychologia.

Hadjikhani, N., Joseph, R., Snyder, J., & Tager-Flusberg, H. (2007). Abnormal activation of the social brain during face perception in autism. Hum Brain Ma.

Happé, F., & Frith, U. (1996). The neuropsychology of autism. Brain.

Happé, F. (1993). Communicative competence and theory of mind in autism: A test of relevance theory. Cognition.

Heerey, E., Keltner, D., & Capps, L. (2003). Making sense of self-conscious emotion: Linking theory of mind and emotion in children with autism. Emotion.

Henry, J. et al.. (2006). Theory of mind following traumatic brain injury: The role of emotion recognition and executive dysfunction. Neuropsychologia.

Hillier, A., & Allinson, L. (2002a). Beyond expectations: Autism, understanding embarrassment, and the relationship with theory of mind. Autism.

Hillier, A., & Allinson, L. (2002b). Understanding embarrassment among those with autism: Breaking down the complex emotion of embarrassment among those with autism. J Autism Dev Disord.

Hoogenhout, M., & Smith, S. M. (2017). Theory of mind predicts severity in autism. Autism.

Hutchins, T., & Prelock, P. (2015). Beyond the theory of mind hypothesis: Using a causal model to understand the nature and treatment of multiple deficits in autism spectrum disorder. Routledge Handbook of Communication Disorders.

Hutchins, T., & Prelock, P. (2016). Technical manual for the theory of mind inventory-2. theoryofmindinventory.com.

Hutchins, T., & Prelock, P. (2017). Theory of mind atlas. theoryofmindinventory.com.

Hutchins, T., Prelock, P., & Bonazinga, L. (2012). Psychometric evaluation of the theory of mind inventory (tomi): A study of typically developing children and children with autism spectrum disorder. J Autism Dev Disord.

Hutchins, T., Prelock, P., & Chase, W. (2008). Test-retest reliability of a theory of mind task battery for children with autism spectrum disorders. Focus Autism Other Dev Disabil.

Ilzarbe, D. et al.. (2020). Neural correlates of theory of mind in autism spectrum disorder, attention-deficit/hyperactivity disorder, and the comorbid condition. Front. Psychiatry.

Jankowski, K., & Takahashi, H. (2014). Cognitive neuroscience of social emotions and implications for psychopathology: Examining embarrassment, guilt, envy, and schadenfreude. Psychiatry Clin Neurosci.

Jensen, J., Helpern, J., Ramani, A., Lu, H., & Kaczynski, K. (2005). Diffusional kurtosis imaging: The quantification of non-gaussian water diffusion by means of magnetic resonance imaging. Magn Reson Med.

Jeurissen, B. et al.. (2014). Multi-tissue constrained spherical deconvolution for improved analysis of multi-shell diffusion mri data. Neuroimage.

Kana, R. et al.. (2015). Aberrant functioning of the theory of mind network in children and adolescents with autism. Molecular Autism volume.

Ke, Z. et al.. (2017). Neural responses to rapid facial expressions of fear and surprise. Frontiers in Psychology.

Konrad, K., Firk, C., & Uhlhaas, P. (2013). Brain development during adolescence: Neuroscientific insights into this developmental period. Deutsches Arzteblatt international.

Koshino, H. et al.. (2008). Fmri investigation of working memory for faces in autism: Visual coding and underconnectivity with frontal areas. Cereb Cortex.

Kuusikko, S. et al.. (2009). Emotion recognition in children and adolescents with autism spectrum disorders. J Autism Dev Disord.

Lacroix, A. et al.. (2014). Facial emotion recognition in 4- to 8-year-olds with autism spectrum disorder: A developmental trajectory approach. Research in Autism Spectrum Disorder.

Lerner, M., Hutchins, T., & Prelock, P. (2011). Brief report: Preliminary evaluation of the theory of mind inventory and its relationship to measures of social skills. J Autism Dev Disord.

Lewis, M. et al.. (1991). Changes in embarrassment as a function of age, sex, and situation. British Journal of Developmental Psychology.

Lord, C., Rutter, M., Dilavore, P., & Risi, S. (1999). Autism diagnostic observation schedule-wps edition. Western Psychological Services.

Mancini, G. et al.. (2018). Recognition of facial emotional expressions among italian pre-adolescents, and their affective reactions. Front. Psychol.

Mandell, D., Listerud, J., & Levy, S. (2002). Race differences in the age at diagnosis among medicaid-eligible children with autism. J Am Acad Child Adolesc Psychiatry.

McCallum, R. (2003). Handbook of nonverbal assessment. SpringerLink.

Modigliani, A., & Blumenfeld, S. (1979). A developmental study of embarrassment and self-presentation. British Journal of Developmental Psychology.

Monk, C. et al.. (2010). Neural circuitry of emotional face processing in autism spectrum disorders. Journal of Psychiatry Neuroscience.

Morrier, M., Hess, K., & Heflin, L. (2008). Ethnic disproportionality in students with autism spectrum disorders. Multicultural Educ Fall.

Müller-Pinzler, L. et al.. (2015). Neural pathways of embarrassment and their modulation by social anxiety. Neuroimage.

Murphy, C. et al.. (2012). Anatomy and aging of the amygdala and hippocampus in autism spectrum disorder: An in vivo magnetic resonance imaging study of asperger syndrome. Autism Res.

National Institute of Mental Health. (2020). The teen brain: 7 things to know.

Nicola, Y. (1984). Young childrens coordination of motive and outcome in judgments of satisfaction and morality. British Journal of Developmental Psychology.

Nomi, J., & Uddin, L. (2015). Face processing in autism spectrum disorders: From brain regions to brain networks. Neuropsychologia.

O’Nions, E. et al.. (2014). Neural bases of theory of mind in children with autism spectrum disorders and children with conduct problems and callous-unemotional traits. Dev Sci.

Pelphrey, K., Morris, J., McCarthy, G., & LaBar, K. (2007). Perception of dynamic changes in facial affect and identity in autism. Soc Cogn Affect Neurosci.

Peterson, C., & Wellman, H. (2005). Steps in theory of mind development for children with deafness or autism. Child Development.

Peterson, C., Wellman, H., & Slaughter, V. (2012). The mind behind the message: Advancing theory of mind scales for typically developing children, and those with deafness, autism, or asperger syndrome. Child Dev.

Phillips, W., Baron-Cohen, S., & Rutter, M. (1995). To what extent can children with autism understand desire? Development and Psychopathology.

Pinkham, A. et al.. (2008). Neural bases for impaired social cognition in schizophrenia and autism spectrum disorders. Schizophr Res.

Poljac, E., Poljac, E., & Wagemans, J. (2012). Reduced accuracy and sensitivity in the perception of emotional facial expressions in individuals with high autism spectrum traits. Autism.

Rieffe, C., Terwogt, M., & Stockmann, L. (2000). Understanding atypical emotions among children with autism. J Autism Dev Disord.

Rieffe, C. et al.. (2010). Preschoolers’ appreciation of uncommon desires and subsequent emotions. British Journal of Developmental Psychology.

Rump, K., Giovannelli, J., Minshew, N., & Strauss, M. (2009). The development of emotion recognition in individuals with autism. Child Development.

Rutter, M. et al.. (2003). Social communication questionnaire. Western Psychological Services.

Scherf, K. S., Luna, B., Minshew, N., & Behrmann, M. (2010). Location, location, location: Alterations in the functional topography of face-but not object- or place-related cortex in adolescents with autism. Front Hum Neurosci.

Schurz, M. et al.. (2017). Specifying the brain anatomy underlying temporo-parietal junction activations for theory of mind: A review using probabilistic atlases from different imaging modalities. Hum Brain Mapp.

Seidner, L., Stipek, D., & Feshbach, N. (1988). A developmental analysis of elementary school-aged children’s concepts of pride and embarrassment. Child Dev.

Setsompop, K. et al.. (2012a). Blipped-controlled aliasing in parallel imaging for simultaneous multislice echo planar imaging with reduced g-factor penalty. Magn Reson Med.

Setsompop, K. et al.. (2012b). Improving diffusion mri using simultaneous multi-slice echo planar imaging. Neuroimage.

Shamay-Tsoory, S. (2008). Recognition of fortune of others emotions in asperger syndrome and high functioning autism. Journal of Autism and Developmental Disorders.

Slaughter, V., & Perez-Zapata, D. (2014). Cultural variations in the development of mind reading. Child Development Perspectives.

Smith, S., & Nichols, T. (2009). Threshold-free cluster enhancement: Addressing problems of smoothing, threshold dependence and localisation in cluster inference. Neuroimage.

Spencer, M. et al.. (2011). A novel functional brain imaging endophenotype of autism: The neural response to facial expression of emotion. Transnational Psychiatry.

Tager-Flusberg, H. (2007). Evaluating the theory of mind hypothesis of autism. Current Directions in Psychological Science.

Takeuchi, M., Harada, M., & Nishitani, H. (2002). Deficiency of “theory of mind” in autism estimated by fmri. International Congress Series.

Tanaka, J., & Sung, A. (2017). The “eye avoidance” hypothesis of autism face processing. J Autism Dev Disord.

Tangney, J., Miller, R., Flicker, L., & Barlow, D. (1996). Are shame, guilt, and embarrassment distinct emotions? Journal of Personality and Social Psychology.

Thabtah, F., & Peebles, D. (2019). A new machine learning model based on induction of rules for autism detection. Health Informatics Journal.

Thirion-Marissiaux, A.-F., & Nader-Grosbois, N. (2007). Theory of mind “emotion”, developmental characteristics and social understanding in children and adolescents with intellectual disabilities. Res Dev Disabil.

Uljarevic, M., & Hamilton, A. (2013). Recognition of emotions in autism: A formal meta-analysis. Journal of Autism and Developmental Disorders.

Volkmar, F. R. et al.. (2014). Handbook of autism and pervasive developmental disorders, fourth edition.

Wallace, S., Coleman, M., & Bailey, A. (2008). An investigation of basic facial expression recognition in autism spectrum disorders. Cognition Emotion.

Weimer, A., Sallquist, J., & Bolnick, R. (2012). Young children’s emotion comprehension and theory of mind understanding. Education and Development.

Wellman, H., Cross, D., & Watson, J. (2011). Meta-analysis of theory of mind development: The truth about false belief. Child Development.

Wellman, H., & Woolley, J. (1990). From simple desires to ordinary beliefs: The early development of everyday psychology. Cognition.

Wellman, H., Fuxi, F., & Petersonn, C. (2011). Sequential progressions in a theory of mind scale: Longitudinal perspectives. Child Development.

Wellman, H., & Liu, D. (2004). Scaling of theory of mind tasks. Child Dev.

Weng, S.-J. et al.. (2011). Neural activation to emotional faces in adolescents with autism spectrum disorders. J Child Psychol Psychiatry.

Wiggins, L., Baio, J., & Rice, C. (2006). Examination of the time between first evaluation and first autism spectrum diagnosis in a population-based sample. J Dev Behav Pediatr.

Williams, D., & Happé, F. (2010). Recognizing social and non-social emotions in self and others: A study of autism. Autism.

Winkler, A. et al.. (2014). Permutation inference for the general linear model. Neuroimage.

Ziv, M., & Frye, D. (2003). The relation between desire and false belief in children’s theory of mind: No satisfaction? Dev Psychol.

